# Bur1-driven G1-to-S phase transition induces hydroxyurea sensitivity in yeast checkpoint mutants

**DOI:** 10.1101/2024.10.21.619526

**Authors:** Stefany Cristine Rodrigues da Silva, Francisco Meirelles Bastos de Oliveira

## Abstract

In *Saccharomyces cerevisiae*, the cyclin-dependent kinase Bur1 is known for its role in promoting transcription elongation and histone modifications, thereby regulating gene expression. In this study, we investigated the genetic interactions between a hypomorphic *BUR1* allele (*bur1-107*) and null mutants of the checkpoint kinases Mec1 and Rad53. Our findings show that the *bur1-107* allele suppresses hydroxyurea (HU) sensitivity in *mec1* and *rad53* mutants, indicating that Bur1 activity may be detrimental in the absence of functional checkpoint signaling. Additionally, the *bur1-107* mutation delays the G1-to-S phase transition, indicating a key role for Bur1 in cell cycle progression. Notably, *bur1-107* reduces γ-H2A accumulation, facilitates S-phase resumption, and supresses the sub-G1 population in HU-treated *mec1* mutants. These findings suggest that Bur1-driven G1-to-S phase progression exacerbates DNA damage and cell death in checkpoint-deficient cells exposed to HU. This study highlights a novel role for Bur1 in modulating the cellular response to replication stress in checkpoint-compromised cells.

## Introduction

During DNA replication, depletion of deoxyribonucleotide triphosphates (dNTPs) can cause replication fork stalling and expose single-stranded DNA (ssDNA), leading to replication stress (RS) (Sogo et al. 2002; Poli et al. 2012). To counteract RS, eukaryotic cells activate the replication stress response (RSR), a set of cellular processes aimed to protect stalled replication forks. Failure to establish an effective RSR can lead to replication fork collapse and increased DNA damage, driving genome instability (Zeman and Cimprich 2014; Saldivar et al. 2017).

In *Saccharomyces cerevisiae*, RSR signaling is mediated by the apical kinase Mec1 and the downstream kinase Rad53, functional homologues of human ATR and Chk1, respectively (Pardo et al. 2017; Lanz et al. 2019; Cussiol et al. 2020). While these kinases were historically known as checkpoint kinases for their role in regulating cell cycle progression (Allen et al. 1994; Weinert et al. 1994), they also perform additional functions. Mec1 and Rad53 suppress late origin firing, stabilize replication forks, and increase dNTP levels by activating ribonucleotide reductase (RNR) genes and degrading the RNR inhibitor Sml1 (Huang et al. 1998; Santocanale and Diffley 1998; Shirahige et al. 1998; Zhao et al. 1998; Lopes et al. 2001; Tercero et al. 2003).

In *S. cerevisiae*, *BUR1* encodes a cyclin-dependent protein kinase homologous to human CDK9 (Yao et al. 2000). Along with its cyclin partner Bur2, Bur1 phosphorylates the C-terminal domain of RNA polymerase II and the transcription elongation factor Spt5, thereby facilitating transcriptional elongation (Murray et al. 2001; Liu et al. 2009). Additionally, Bur1 is involved in co-transcriptional histone modifications, including the monoubiquitination of histone H2B and the mono- and trimethylation of histone H3 (Laribee et al. 2005; Wood et al. 2005; Chu et al. 2006). Through these mechanisms, the Bur1-Bur2 complex regulates the expression of a broad range of genes (Laribee et al. 2005).

Growing evidence suggests a functional interplay between the Bur1-Bur2 complex and the RSR pathway. Bur1 was found to physically interact with Rfa1, a subunit of the replication protein A (RPA) complex and a key sensor of RS (Clausing et al. 2010). Bur2-deficient cells exhibits sensitivity to hydroxyurea (HU), an RNR inhibitor that depletes dNTPs and is commonly used to induce RS (Lau et al. 2010). Additionally, HU-treated *bur1* mutants display an accumulation of Rad52 foci, suggesting that Bur1 is involved in preventing the colpase of stalled replication forks (Clausing et al. 2010).

To further explore Bur1’s role in the RSR we conducted a genetic interaction analysis by combining a hypomorphic mutant of *BUR1* (*bur1-107*) with null alleles of *MEC1* or *RAD53*. Our study shows that the *bur1-107* mutantion alleviates the HU sensitivity in *mec1* and *rad53* mutants. Additionally, we found that the *bur1-107* mutant delays the transition from G1 to S-phase. Notably, *bur1-107* reduces γ-H2A accumulation, facilitates S-phase resumption and supresses the sub-G1 population in HU-treated *mec1*Δ cells. These findings suggest that, in the absence of a functional checkpoint, Bur1 may have a detrimental effect, potentially by driving an inappropriate transition from G1 to S-phase.

## Materials and methods

### Strains and plasmids

Unless otherwise noted, all strains are derived from the S288C background. Gene deletions were achieved by replacing the endogenous allele with drug-resistance cassettes or auxotrophic markers, generated by PCR (Longtine et al. 1998). Yeast transformations were performed using the lithium acetate method (Gietz 2014), and successful knockouts were confirmed by PCR with specific primers. To construct the *bur1-107* strain, the *BUR1* open reading frame and a 50 bp upstream region were cloned into the pFA6a-kanMX6 vector using *PacI* and *AscI*. Site-directed mutagenesis introduced the G52R and G222N mutations. A 466 bp downstream region of *BUR1* was then cloned into the plasmid using *PmeI* and *SpeI*. The *bur1-107* construct was excised with *PacI* and *SpeI* and integrated into the *BUR1* locus in a diploid strain. After sporulation, haploid strains carrying the *bur1-107* allele were isolated by tetrad dissection and confirmed by PCR and kanMX6 marker selection. The same procedure was used to generate isogenic strains with the wild-type *BUR1* allele.

### Cell growth, media, and synchronization

Unless stated otherwise, cells were cultured in YPD medium (1% yeast extract, 2% peptone, 2% dextrose) at 30°C to logarithmic phase. For synchronization, *bar1*Δ cells were treated with α-factor (50 ng/mL for *bar1*Δ strains, 5 µg/mL for *bar1*Δ *mec1*Δ or *bar1*Δ *rad53*Δ strains) for 2,5 hours (Amberg et al. 2006). Cells were then released from G1 arrest by centrifugation and resuspended in fresh medium containing 50 µg/mL pronase, with or without HU. The specific HU concentrations used are indicated in the corresponding figures and figure legends.

### Drug sensitivity assays

Overnight cultures were inoculated in YPD medium to an initial OD_600_ of 0.1 and grown to mid-log phase. Cultures were then adjusted to an OD_600_ of 1.0, and five-fold serial dilutions were spotted onto plates with or without HU. Plates were incubated at 30°C for 1–3 days, then inspected for growth and photographed. For drug sensitivity assays with 6-azauracil (6AU), pRS316-bearing cells were cultured in synthetic complete medium lacking uracil (SC-URA). The specific HU and 6AU concentrations used are indicated in the corresponding figures and figure legends.

### Flow cytometry

A total of 5 x 10 yeast cells were harvested, washed with distilled water, and sonicated (10 seconds at 20% amplitude) to disperse clumps. Cells were fixed in 1 mL of 70% ethanol overnight at 4°C. After fixation, cells were centrifuged, ethanol removed, and pellets resuspended in 50 mM sodium citrate buffer (pH 7.2). The samples were treated with 250 µg/mL RNase A at 37°C overnight, followed by 2 hours with 500 µg/mL proteinase K at 50°C. SYTOX™ Green (0.25 µL; Thermo Fisher Scientific) was then added, and samples were incubated for 1 hour at room temperature in the dark. Data acquisition was performed using a BD FACSCalibur™ flow cytometer.

### Western blotting

Yeast cultures were grown to mid-log phase and treated as described in the figure legends. Cells were harvested and washed with TE buffer (pH 8.0) containing 1 mM PMSF. Pellets were lysed by bead beating with 0.5-mm glass beads for three 5-minute cycles (1-minute rest between cycles) at 4°C in lysis buffer (150 mM NaCl, 50 mM Tris pH 8.0, 5 mM EDTA, 0.2% NP-40) with cOmplete^TM^ EDTA-free protease inhibitor cocktail (Roche), PhosStop^TM^ phosphatase inhibitor cocktail (Roche), and 1 mM PMSF. Lysates were boiled in Laemmli buffer and subjected to SDS-PAGE. Proteins were transferred to PVDF membranes by wet transfer and incubated with specific primary antibodies: anti-γ-H2A (ab15083, Abcam, 1:1,000), anti-Rad53 (ab166859, Abcam, 1:5,000), and anti-PGK1 (459250, Invitrogen, 1:5,000). After primary antibody incubation, membranes were treated with ECL HRP-linked secondary antibodies (anti-mouse NA931, Cytiva, 1:5,000; anti-rabbit NA934, Cytiva, 1:5,000). Blots were developed with ECL^TM^ Prime reagent (Amersham) and imaged using an Amershan^TM^ Imager 600.

## Results

### Bur1 exacerbates HU sensitivity in *mec1* and *rad53* mutants

To further investigate Bur1’s role in RSR, we conducted a pairwise genetic interaction analysis between *BUR1* and the checkpoint kinase genes *MEC1* and *RAD53*. Given that *BUR1* is essential, we used the hypomorphic *bur1-107* mutant, which contains two amino acid substitutions: G52R at the N-terminus and G222N within the activation segment of the kinase domain (Fig. 1a) (Clausing et al. 2010).

**Figure 1.**
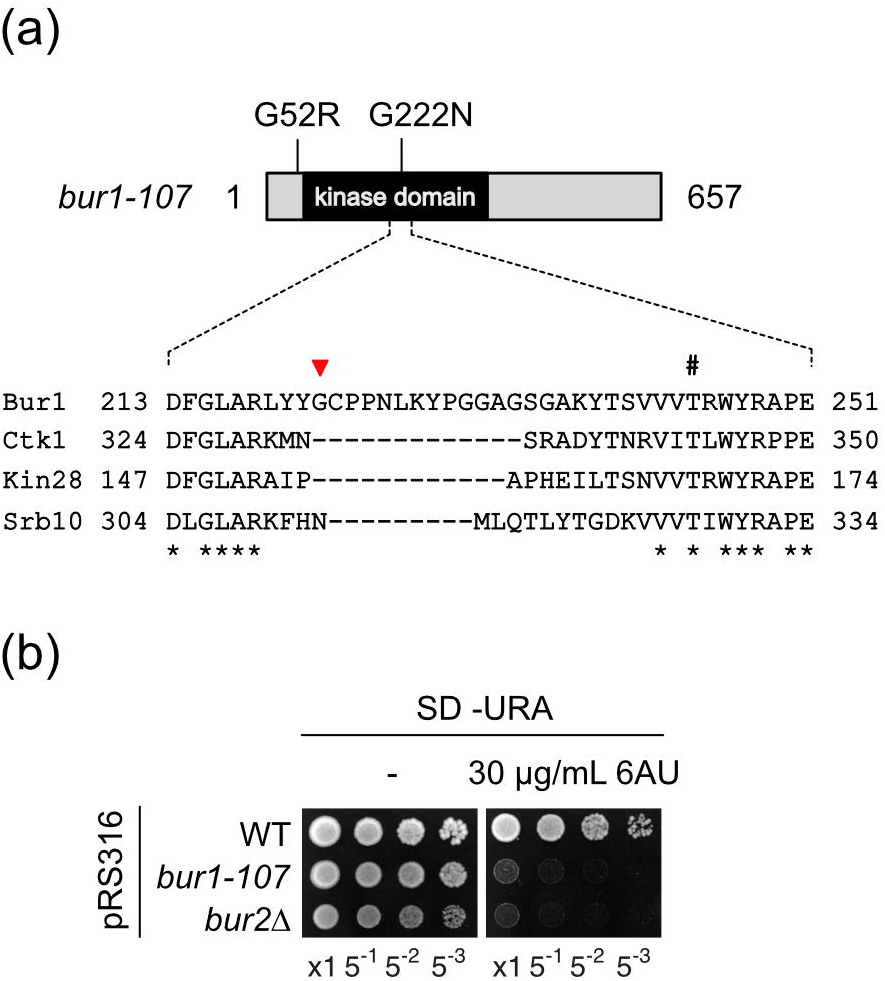
*bur1-107* and *bur2* mutants are sensitive to 6AU. (a) Schematic of the *bur1-107* protein, highlighting the kinase domain and the G52R (N-terminal) and G222N (kinase domain) mutations. The dotted line represents the amino acid alignment of the activation segments between yeast transcription-related Cdks Bur1, Ctk1, Kin28 and Srb10, spanning from the conserved DFG motif to the APE motif. The hashtag marks the conserved phosphorylation-dependent threonine residue in the T-loop (Malumbres 2014). The red arrow indicates the G222 residue mutated in *bur1-107*. Numbers indicate amino acid positions, with asterisks marking conserved residues. (b) Serial 1:5 dilutions of pRS316-bearing strains with the indicated genotypes were spotted onto SC-URA plates with or without 6-azauracil (6AU). Plates were incubated at 30°C for 3 days.

The *bur1-107* mutant showed increased sensitivity to 6-azauracil (6AU), an inhibitor of GTP and UTP synthesis (Fig. 1b). While reduced GTP and UTP levels are not lethal to wild-type cells, they significantly increase sensitivity in conjunction with mutations that impair transcriptional elongation (Exinger and Lacroute 1992). Additionally, we confirmed that the *bur2*Δ mutant is also sensitive to 6AU, further supporting the role of the Bur1-Bur2 complex in facilitating transcription elongation (Fig. 1b).

Since *MEC1* and *RAD53* are essential, the *mec1*Δ and *rad53*Δ mutants used in the genetic interaction assays were generated in a *sml1*Δ background (Zhao et al. 1998). Notably, the *bur1-107* mutant partially suppressed the sensitivity of both *mec1*Δ *sml1*Δ and *rad53*Δ *sml1*Δ cells to low doses of HU (Fig. 2a). Similarly, the deletion of *BUR2* also alleviated HU sensitivity in *mec1*Δ *sml1*Δ cells (Supplementary Fig. S1). These findings suggest that, in the absence of a functional checkpoint signaling, the Bur1-Bur2 complex exacerbates the detrimental effects of HU treatment.

**Figure 2.**
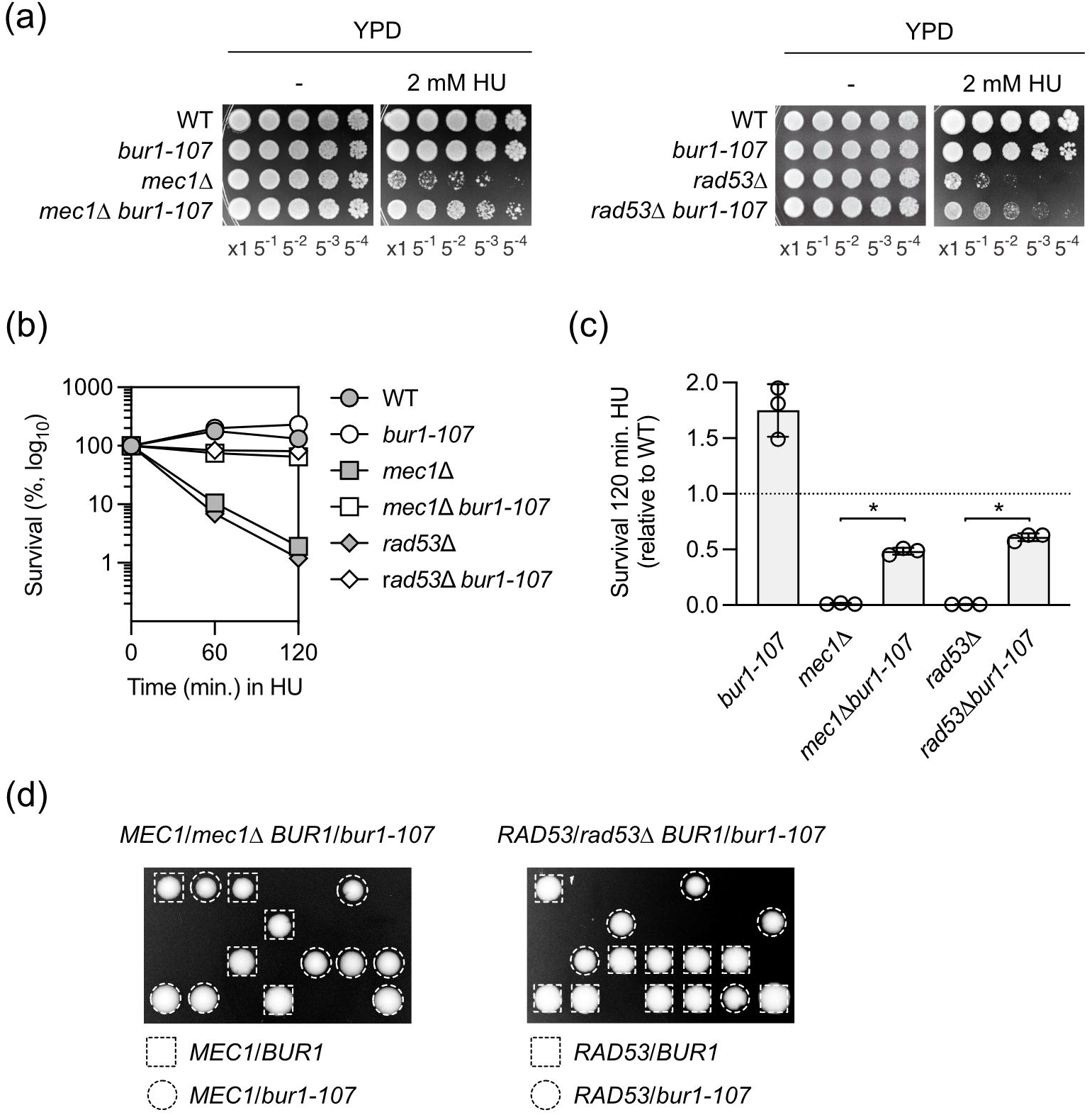
*bur1-107* improves viability of *mec1* and *rad53* mutants treated with HU. (a) and (b) Serial dilutions (1:5) of strains with indicated genotypes were spotted onto YPD plates with or without hydroxyurea (HU). Plates were incubated at 30°C for 3 days. (b) Strains with indicated genotypes were synchronized in G1 with α-factor, released into YPD with pronase and 200 mM HU, and assessed for survival at indicated time points by colony-forming units on fresh YPD plates. Plates were incubated at 30°C for 3 days. Data are the mean of at least three experiments. (d) Relative survival of S-phase cells treated with 200 mM HU for 2 hours, compared to WT control. Data represent the mean ± SEM from at least three independent experiments. *P < 0.05 (Student’s unpaired t-test). All strains used in the experiments shown in panels (a) to (d) carry the *sml1*Δ mutation. (e) Meiotic tetrads from *MEC1*/*mec1*Δ *BUR1*/*bur1-107* or *RAD53*/*rad53*Δ *BUR1*/*bur1-107* diploids were dissected on YPD and incubated at 30°C for 3 days. WT and *bur1-107* single-mutant spores are indicated by squares and circles, respectively.

To evaluate the impact of the *bur1-107* mutation on the viability of *mec1*Δ *sml1*Δ or *rad53*Δ *sml1*Δ mutants exposed to HU during a single round of DNA replication, cells were synchronized in G1 phase using α-factor and subsequently released into S-phase in the presence of 200 mM HU for two hours. As shown in Fig. 2b and c, *bur1-107* partially restored viability in *mec1*Δ *sml1*Δ and *rad53*Δ *sml1*Δ mutants after two hours of HU exposure.

Next, we questioned whether *bur1-107* would be sufficient to bypass the essential functions of *MEC1* or *RAD53*. Diploid strains heterozygous for *bur1-107* and either *mec1* or *rad53* alleles were generated, and spore viability was assessed following tetrad dissection. As expected, *mec1*Δ and *rad53*Δ spores were inviable and failed to form colonies (Fig. 2d). Although *bur1-107* spores grew to nearly wild-type size, no colony formation was observed for *bur1-107 mec1*Δ or *bur1-107 rad53*Δ spores. These results suggest that, unlike *sml1*Δ, *bur1-107* cannot compensate for the loss of *MEC1* or *RAD53*.

### *bur1-107* delays the transition from G1 to S-phase independently of Mec1

Throughout this study, we noted that the *bur1-107* mutant exhibited slower growth compared to wild-type cells (Supplementary Fig. S2). This is consistent with previous findings showing that Bur1-deficient cells undergo an extended G1 phase (Jin et al. 2022). To assess whether the *bur1-107* mutant is defective in the G1-to-S phase transition, cells were synchronized in G1, released into S-phase, and DNA content was analyzed by flow cytometry (Fig. 3a). To prevent complete DNA replication stalling, experiments were conducted with 20 mM of HU (Fig. 3c). As expected, *mec1*Δ *sml1*Δ cells progressed through S phase slightly faster than wild-type cells (*sml1*Δ). However, despite their accelerated progression, *mec1*Δ *sml1*Δ cells were unable to complete DNA replication, even 120 minutes after release from α-factor arrest. Interestingly, *bur1-107 sml1*Δ and *bur1-107 mec1*Δ *sml1*Δ cells exhibited a more pronounced delay in G1/S progression compared to *mec1*Δ *sml1*Δ cells, both with or without HU (Fig. 3b and c). Consistent with its role in promoting the G1/S transition, *bur1-107 sml1*Δ and *bur1-107 mec1*Δ *sml1*Δ mutants showed a delay in bud formation compared to *mec1*Δ *sml1*Δ (Fig. 3d). These data suggest that Bur1 is required for cell cycle progression from G1 to S-phase, independently of Mec1.

**Figure 3.**
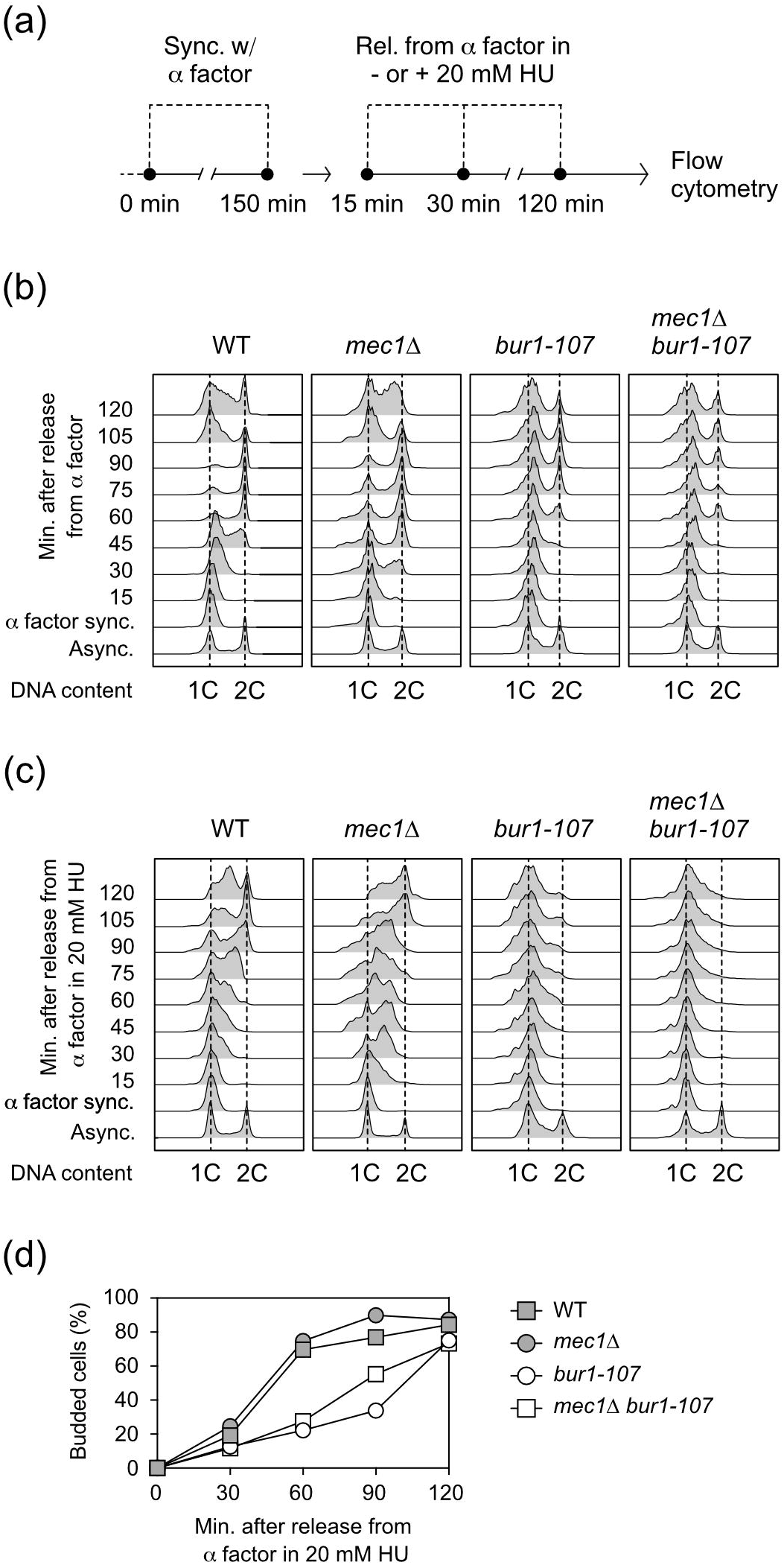
*bur1-107* delays the progression from G1 to S-phase. (a) Exponentially growing asynchronous cells were arrested in G1 with α-factor for 2,5 hours, then released into YPD at 25°C, either (b) without or (c) with 20 mM HU. DNA content was analyzed by flow cytometry at the indicated time points. (d) Exponentially growing strains with the indicated genotypes were arrested in G1 with α-factor for 2,5 hours and then released into YPD with pronase and 20 mM HU at 25°C. Budding index was measured at the specified time points to monitor cell cycle progression. All strains carry the *sml1*Δ mutation.

### BUR1 increases DNA damage and impairs the resumption of S-phase in mec1 mutants treated with HU

To further investigate the effect of Bur1 in *mec1*Δ *sml1*Δ cells, we evaluated the impact of the *bur1-107* mutation on DNA damage signaling. Cells were synchronized in G1 and then released into S-phase with 200 mM of HU for up to 120 minutes (Fig. 4a, b and Supplementary Fig. S3). Consistent with previous studies, Rad53 activation was observed at 30 and 60 minutes in wild-type cells (*sml1*Δ), while no increase in γ-H2A signal was detected (Puddu et al. 2011). In contrast, *mec1*Δ *sml1*Δ cells showed γ-H2A accumulation and reduced Rad53 activation. Intriguingly, *bur1-107 mec1*Δ *sml1*Δ cells displayed a significant decrease in Rad53 phosphorylation and γ-H2A signal compared to *mec1*Δ *sml1*Δ cells. These findings demonstrate that, in *mec1*Δ *sml1*Δ cells exposed to HU, Bur1 exacerbates RS and promotes DNA damage.

**Figure 4.**
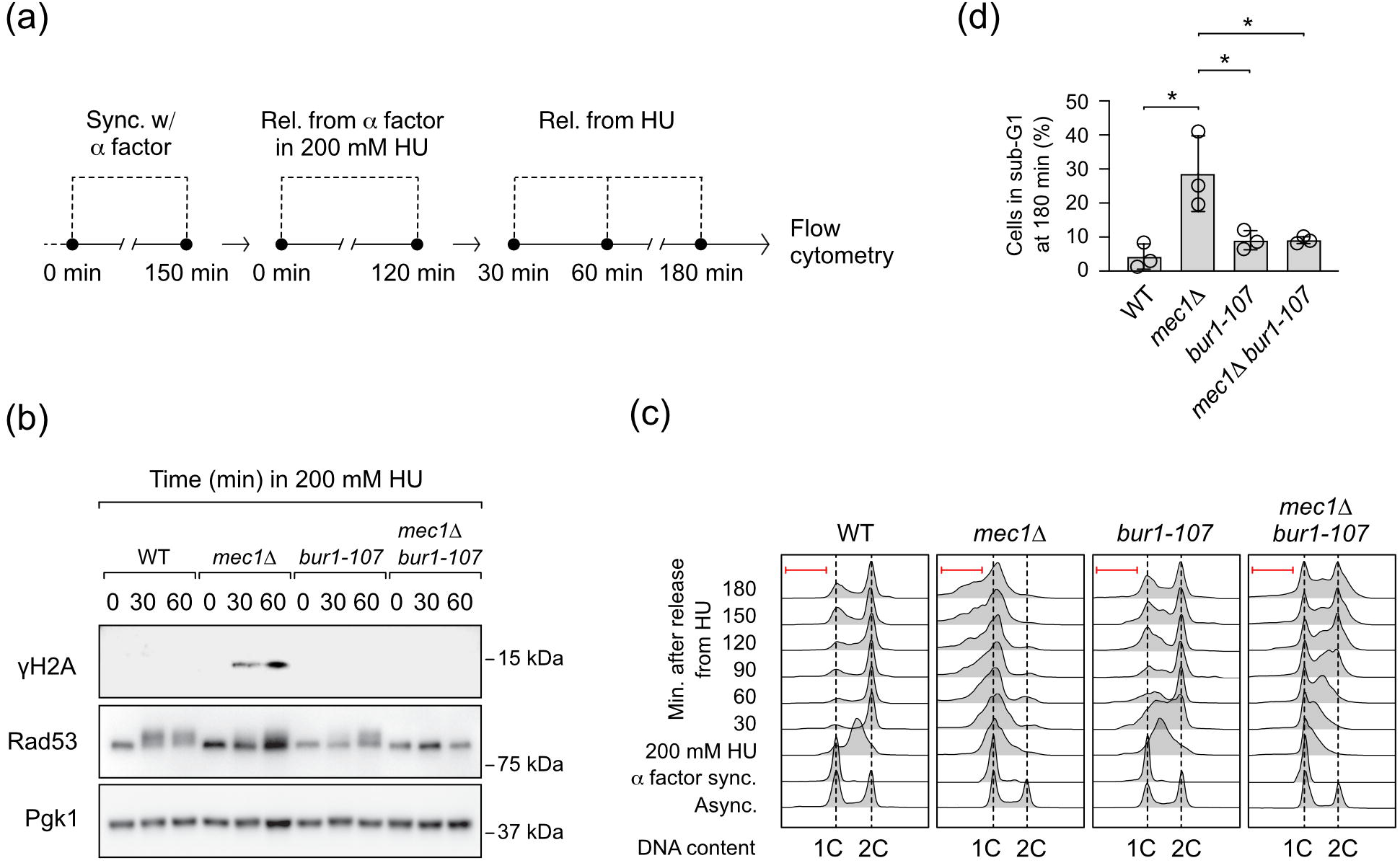
*bur1-107* supresses DNA damage signaling and facilitates the resumption of S-phase in HU-treated *mec1* mutants. (a) Exponentially growing asynchronous cells were arrested in G1 phase with α-factor for 2,5 hours, followed by release into YPD containing 200 mM HU for an additional 2 hours. After HU treatment, cells were released into fresh YPD and incubated for 3 hours at 30°C. (b) Samples were collected at the specified time points post-release from α-factor for Western blot analysis to detect γ-H2A and Rad53. (c) DNA content was analyzed by flow cytometry at the indicated time points. Red lines indicate sub-G1 populations. (d) Percentage of cells in sub-G1 phase at 180 minutes post-HU release, relative to total cell cycle phases. Data represent the mean ± SEM from at least three independent experiments. *P < 0.05 (Student’s unpaired t-test). All strains carry the *sml1*Δ mutation.

To determine whether Bur1 influences the resumption of S-phase in HU-treated *mec1*Δ *sml1*Δ cells, we analyzed cell cycle progression following a two-hour HU-induced replication block (Fig. 4a and c). *bur1-107 sml1*Δ cells initially exhibit a delay in S-phase progression 30 minutes after HU removal but resumed replication between 90 and 120 minutes. In contrast, *mec1*Δ *sml1*Δ cells showed a pronounced defect in replication resumption, failing to reach the 2C DNA content peak even 180 minutes post-HU removal. Consistent with the observed lethality in *mec1*Δ *sml1*Δ cells treated with HU (Fig. 2c and d), a sub-G1 population was detected at 180 minutes after HU removal (Fig. 4c, red brackets). Interestingly, although *bur1-107 mec1*Δ *sml1*Δ mutants progressed slowly through S-phase, a subset of these cells reached the 2C DNA content peak at 120 minutes post-HU release. Notably, by 180 minutes after HU removal, the *bur1-107 mec1*Δ *sml1*Δ mutant exhibited a lower percentage of cells in sub-G1 compared to *mec1*Δ *sml1*Δ cells (Fig. 4d). These findings suggest that Bur1 exacerbates DNA replication defects in *mec1*Δ *sml1*Δ cells following HU treatment, resulting in increased DNA damage and cell death.

## Discussion

Growing evidence suggests a functional interplay between the Bur1-Bur2 complex and the RSR pathway. Our study reveals a genetic interaction between *BUR1* and the genes encoding the checkpoint signaling kinases Mec1 and Rad53. Notably, we observed that the *bur1-107* mutant alleviates HU sensitivity in *mec1*Δ *sml1*Δ and *rad53*Δ *sml1*Δ cells. These findings imply that in the absence of functional checkpoint signaling, Bur1 may have a detrimental impact on cells treated with HU.

Bur1 was initially identified as a CDC28/cdc2-related kinase essencial for the adaptative response to pheromone-induced G1 arrest (Irie et al. 1991). Although the precise mechanism is not fully understood, it has been proposed that Bur1 aids the transition from G1 phase by activating G1 cyclins. Supporting this hypothesis, Bur1-deficient cells show increased retention of nuclear Whi5, a known G1 marker (Jin et al. 2022). Consistently, we observed that the *bur1-107* mutant exhibits a prolonged doubling time and delayed progression from G1 to S-phase.

The Bur1-Bur2 complex is critical for facilitating transcription elongation, thereby regulating the expression of a wide range of genes. In particular, the *bur1-107* mutation impairs transcription elongation and results in decreased mRNA levels of cyclins Cln2, Clb1, and Clb6 (Clausing et al. 2010). This suggests that *bur1-107* might also impact the expression of other genes essencial for cell cycle regulation, particularly those involved in the G1 to S-phase transition. Additionaly, It is possible that Bur1 directly phosphorylates substrates that promote the G1 to S-phase transition, independently of its role in transcriptional regulation.

Our findings reveal that the *bur1-107* mutation supresses the lethality of *mec1*Δ *sml1*Δ cells and facilitates the resumption of S-phase following a two-hour replication block with HU. Adittionally, the *bur1-107* mutation significantly reduces the accumulation of γ-H2A in HU-treated *mec1*Δ *sml1*Δ cells. It is reasonable to hypothesize that by delaying the G1 to S-phase transition, the *bur1-107* mutation alleviates the detrimental effects of HU-induced RS in *mec1*Δ *sml1*Δ cells. Indeed, the observed lethality in *mec1* and *rad53* mutants is largely attributed to a Cdc28-dependent function during G1 phase.

Consistently, Cdc28-defective strains supress the HU-induced lethality in *mec1*Δ *sml1*Δ and *rad53*Δ *sml1*Δ cells (Manfrini et al. 2012). Similarly, a *cln2*Δ mutant, which delays the G1-to-S phase transition, also suppresses the requirement for Mec1 and Rad53 during an unperturbed cell cycle (Manfrini et al. 2012). Although the precise mechanism of suppression remains unclear, it has been proposed that delaying the G1-to-S phase transition allows *mec1* and *rad53* mutants additional time to accumulate sufficient dNTPs for DNA replication, thereby alleviating RS (Vallen and Cross 1999). Our data support this model, showing that in *mec1*Δ cells, the downregulation of *BUR1*, in combination with *SML1* inactivation, suppresses HU sensitivity, likely by increasing dNTP pools and thus reducing the dependence on Mec1 for checkpoint activation.

Interestingly, while the Bur1-Bur2 complex seems to exacerbate stress in checkpoint kinase-deficient strains treated with low doses of HU, the *bur1-107* and *bur2*Δ mutants exhibit increased sensitivity to higher HU concentrations (Supplementary Fig. S4). Consistent with this observation, *bur1-107* exhibit an increase in Rad52 foci in HU-treated cells, suggesting that the Bur1-Bur2 complex may regulate additional processes crucial for protecting cells from the detrimental effects of prolonged exposure to high HU doses (Clausing et al. 2010).

In human cells, the Bur1 homologue, CDK9, is crucial for maintaining genome stability. Depletion of CDK9 impairs cell cycle recovery in response to RS and induces spontaneous DNA damage during replication (Yu et al. 2010). Additionaly, CDK9-deficient cells exhibit altered γ-H2AX foci dynamics and reduced efficiency in homologous recombination repair following ionizing radiation (Nepomuceno et al. 2017).

Here, we demonstrate that despite evidence supporting a protective role for Bur1 during RS, it can exert a detrimental effect in cells with compromised cell cycle checkpoints. Specifically, Bur1-driven G1-to-S phase transition exacerbates RS and promotes cell death in *mec1* mutants exposed to HU. To fully elucidate this dual role of Bur1, future studies should focus on systematically identifying its transcriptional targets and phosphorylation substrates, along with investigating their precise roles in regulating cell cycle progression and the cellular response to RS.

## Data availability

Strains and plasmids used in this study are available upon request. Supplementary Tables 1 and 2 provide detailed lists of the yeast strains and plasmids utilized. Supplementary Figures S1 and S2 include additional data supporting Figures 2 and 3, respectively, while Supplementary Figures S3 and S4 offer supplementary data supporting Figure 4 and the discussion section, respectively.

## Acknowledgments

We thank Marcelo Fantappié, Ronaldo Mohana and Marcelo de Pádula for providing access to their laboratory and equipment. We also thank Claudio Masuda, José Renato Cussiol, Marcelo Alex de Carvalho and Thales Nepomuceno for their valuable comments and suggestions and Patricia Abrão Possik and Marcus Smolka for sharing reagents. Our appreciation goes to Anderson Amarante and Karina Lani Silva for their technical support. Additionally, we acknowledge the Plataforma de Sequenciamento de DNA (PSEQDNA-UFRJ) for DNA sequencing and the Plataforma de Imuno Análise (PIA-UFRJ) for flow cytometry analysis of DNA content.

## Authors contribution

FMBdO conceived the project, FMBdO and SCRdS wrote the paper, designed the experiments and analyzed the data. SCRdS performed the experiments.

---

Figure S1, related to Figure 2. *bur2***Δ** suppresses sensitivity in *mec1* mutants treated with HU.

Serial dilutions (1:5) of strains with indicated genotypes were spotted onto YPD plates with or without hydroxyurea (HU). Plates were incubated at 30°C for 3 days. All strains used carry the *sml1*Δ mutation.

Figure S2, related to Figure 3. *bur1-107* exhibits slower growth compared to WT cells.

Extended growth curves of the indicated strains cultured in YPD medium. The gray shaded area highlights the section of the curve used to calculate doubling time. Data represent the mean of at least six replicate cultures.

Figure S3, related to Figure 4. *bur1-107* suppresses γ-H2A signaling in HU-treated *mec1* mutants between 90 and 120 minutes post-release from α-factor.

Exponentially growing cultures of strains with the indicated genotypes were arrested in G1 with α-factor for 2,5 hours, then released into YPD with pronase and 200 mM HU at 30°C. At the indicated times post-release, cell samples were collected for Western blot analysis to detect γ-H2A. All strains used carry the *sml1*Δ mutation.

Figure S4, related to discussion. *bur1-107* and *bur2* mutants exhibit sensitivity to prolonged exposure to high doses of hydroxyurea.

Exponentially growing cultures of strains with the indicated genotypes were serially diluted (1:5) and spotted onto YPD plates with or without hydroxyurea (HU) at the indicated concentrations. The plates were incubated at 30°C for 3 days. All strains carry the *sml1*Δ mutation.

## References

Allen JB, Zhou Z, Siede W, Friedberg EC, Elledge SJ. 1994. The SAD1/RAD53 protein kinase controls multiple checkpoints and DNA damage-induced transcription in yeast. Genes Dev. 8(20):2401–2415. doi:10.1101/gad.8.20.2401.

Amberg DC, Burke DJ, Strathern JN. 2006. Inducing Yeast Cell Synchrony: α-Factor Arrest Using *bar1* Mutants. Cold Spring Harb Protoc. 2006(1):pdb.prot4173. doi:10.1101/pdb.prot4173.

Chu Y, Sutton A, Sternglanz R, Prelich G. 2006. The Bur1 Cyclin-Dependent Protein Kinase Is Required for the Normal Pattern of Histone Methylation by Set2. Mol Cell Biol. 26(8):3029–3038. doi:10.1128/mcb.26.8.3029-3038.2006.

Clausing E, Mayer A, Chanarat S, Müller B, Germann SM, Cramer P, Lisby M, Strässer K. 2010. The transcription elongation factor Bur1-Bur2 interacts with replication protein A and maintains genome stability during replication stress. Journal of Biological Chemistry. 285(53):41665–41674. doi:10.1074/jbc.M110.193292.

Cussiol JRR, Soares BL, Oliveira FMB de. 2020. From yeast to humans: Understanding the biology of DNA Damage Response (DDR) kinases. Genet Mol Biol. 43(1 suppl 1). doi:10.1590/1678-4685-gmb-2019-0071.

Exinger F, Lacroute F. 1992. 6-Azauracil inhibition of GTP biosynthesis in Saccharomyces cerevisiae. Curr Genet. 22(1):9–11. doi:10.1007/BF00351735.

Gietz RD. 2014. Yeast Transformation by the LiAc/SS Carrier DNA/PEG Method. p. 1–12.

Huang M, Zhou Z, Elledge SJ. 1998. The DNA Replication and Damage Checkpoint Pathways Induce Transcription by Inhibition of the Crt1 Repressor. Cell. 94(5):595–605. doi:10.1016/S0092-8674(00)81601-3.

Irie K, Nomoto S, Miyajima I, Matsumoto K. 1991. SGV1 encodes a CDC28/cdc2-related kinase required for a Gα subunit-mediated adaptive response to pheromone in S. cerevisiae. Cell. 65(5):785–795. doi:10.1016/0092-8674(91)90386-D.

Jin Y, Jin N, Oikawa Y, Benyair R, Koizumi M, Wilson TE, Ohsumi Y, Weisman LS. 2022. Bur1 functions with TORC1 for vacuole-mediated cell cycle progression. EMBO Rep. 23(4). doi:10.15252/embr.202153477.

Lanz MC, Dibitetto D, Smolka MB. 2019. DNA damage kinase signaling: checkpoint and repair at 30 years. EMBO J. 38(18):e101801. doi:10.15252/embj.2019101801 PMID - 31393028.

Laribee RN, Krogan NJ, Xiao T, Shibata Y, Hughes TR, Greenblatt JF, Strahl BD. 2005. BUR kinase selectively regulates H3 K4 trimethylation and H2B ubiquitylation through recruitment of the PAF elongation complex. Current Biology. 15(16):1487–1493. doi:10.1016/j.cub.2005.07.028.

Lau NC, Mulder KW, Brenkman AB, Mohammed S, van den Broek NJF, Heck AJR, Timmers HTM. 2010. Phosphorylation of Not4p functions parallel to BUR2 to regulate resistance to cellular stresses in saccharomyces cerevisiae. PLoS One. 5(4). doi:10.1371/journal.pone.0009864.

Liu Y, Warfield L, Zhang C, Luo J, Allen J, Lang WH, Ranish J, Shokat KM, Hahn S. 2009. Phosphorylation of the Transcription Elongation Factor Spt5 by Yeast Bur1 Kinase Stimulates Recruitment of the PAF Complex. Mol Cell Biol. 29(17):4852–4863. doi:10.1128/mcb.00609-09.

Longtine MS, McKenzie A, Demarini DJ, Shah NG, Wach A, Brachat A, Philippsen P, Pringle JR. 1998. Additional modules for versatile and economical PCR-based gene deletion and modification in Saccharomyces cerevisiae. Yeast. 14(10):953–961. doi:10.1002/(SICI)1097-0061(199807)14:10<953::AID-YEA293>3.0.CO;2-U.

Lopes M, Cotta-Ramusino C, Pellicioli A, Liberi G, Plevani P, Muzi-Falconi M, Newlon CS, Foiani M. 2001. The DNA replication checkpoint response stabilizes stalled replication forks. Nature. 412(6846):557–561. doi:10.1038/35087613.

Malumbres M. 2014. Cyclin-dependent kinases. Genome Biol. 15(6):122. doi:10.1186/gb4184.

Manfrini N, Gobbini E, Baldo V, Trovesi C, Lucchini G, Longhese MP. 2012. G 1 /S and G 2 /M Cyclin-Dependent Kinase Activities Commit Cells to Death in the Absence of the S-Phase Checkpoint. Mol Cell Biol. 32(24):4971–4985. doi:10.1128/mcb.00956-12.

Murray S, Udupa R, Yao S, Hartzog G, Prelich G. 2001. Phosphorylation of the RNA Polymerase II Carboxy-Terminal Domain by the Bur1 Cyclin-Dependent Kinase. Mol Cell Biol. 21(13):4089–4096. doi:10.1128/mcb.21.13.4089-4096.2001.

Nepomuceno TC, Fernandes VC, Gomes TT, Carvalho RS, Suarez-Kurtz G, Monteiro AN, Carvalho MA. 2017. BRCA1 recruitment to damaged DNA sites is dependent on CDK9. Cell Cycle. 16(7):665–672. doi:10.1080/15384101.2017.1295177.

Pardo B, Crabbé L, Pasero P. 2017. Signaling pathways of replication stress in yeast. FEMS Yeast Res. 17(2). doi:10.1093/femsyr/fow101.

Poli J, Tsaponina O, Crabbé L, Keszthelyi A, Pantesco V, Chabes A, Lengronne A, Pasero P. 2012. dNTP pools determine fork progression and origin usage under replication stress. EMBO J. 31(4):883–894. doi:10.1038/emboj.2011.470.

Puddu F, Piergiovanni G, Plevani P, Muzi-Falconi M. 2011. Sensing of Replication Stress and Mec1 Activation Act through Two Independent Pathways Involving the 9-1-1 Complex and DNA Polymerase ε. PLoS Genet. 7(3):e1002022. doi:10.1371/journal.pgen.1002022.

Saldivar JC, Cortez D, Cimprich KA. 2017. The essential kinase ATR: ensuring faithful duplication of a challenging genome. Nat Rev Mol Cell Biol. 18(10):622–636. doi:10.1038/nrm.2017.67.

Santocanale C, Diffley JFX. 1998. A Mec1- and Rad53-dependent checkpoint controls late-firing origins of DNA replication. Nature. 395(6702):615–618. doi:10.1038/27001.

Shirahige K, Hori Y, Shiraishi K, Yamashita M, Takahashi K, Obuse C, Tsurimoto T, Yoshikawa H. 1998. Regulation of DNA-replication origins during cell-cycle progression. Nature. 395(6702):618–621. doi:10.1038/27007.

Sogo JM, Lopes M, Foiani M. 2002. Fork Reversal and ssDNA Accumulation at Stalled Replication Forks Owing to Checkpoint Defects. Science (1979). 297(5581):599–602. doi:10.1126/science.1074023.

Tercero JA, Longhese MP, Diffley JFX. 2003. A Central Role for DNA Replication Forks in Checkpoint Activation and Response. Mol Cell. 11(5):1323–1336. doi:10.1016/S1097-2765(03)00169-2.

Vallen EA, Cross FR. 1999. Interaction Between the MEC1-Dependent DNA Synthesis Checkpoint and G1 Cyclin Function in Saccharomyces cerevisiae. Genetics. 151(2):459–471. doi:10.1093/genetics/151.2.459.

Weinert TA, Kiser GL, Hartwell LH. 1994. Mitotic checkpoint genes in budding yeast and the dependence of mitosis on DNA replication and repair. Genes Dev. 8. doi:10.1101/gad.8.6.652.

Wood A, Schneider J, Dover J, Johnston M, Shilatifard A. 2005. The Bur1/Bur2 complex is required for histone H2B monoubiquitination by Rad6/Bre1 and histone methylation by COMPASS. Mol Cell. 20(4):589–599. doi:10.1016/j.molcel.2005.09.010.

Yao S, Neiman A, Prelich G. 2000. *BUR1* and *BUR2* Encode a Divergent Cyclin-Dependent Kinase–Cyclin Complex Important for Transcription In Vivo. Mol Cell Biol. 20(19):7080–7087. doi:10.1128/MCB.20.19.7080-7087.2000.

Yu DS, Zhao R, Hsu EL, Cayer J, Ye F, Guo Y, Shyr Y, Cortez D. 2010. Cyclin-dependent kinase 9-cyclin K functions in the replication stress response. EMBO Rep. 11(11):876–882. doi:10.1038/embor.2010.153.

Zeman MK, Cimprich KA. 2014. Causes and consequences of replication stress. Nat Cell Biol. 16(1):2–9. doi:10.1038/ncb2897 PMID - 24366029.

Zhao X, Muller EGD, Rothstein R. 1998. A Suppressor of Two Essential Checkpoint Genes Identifies a Novel Protein that Negatively Affects dNTP Pools. Mol Cell. 2(3):329–340. doi:10.1016/S1097-2765(00)80277-4.

