## Supplementary figures and images for "Bur1-driven G1-to-S phase transition induces hydroxyurea sensitivity in yeast checkpoint mutants"

### Supplemental Figure 1

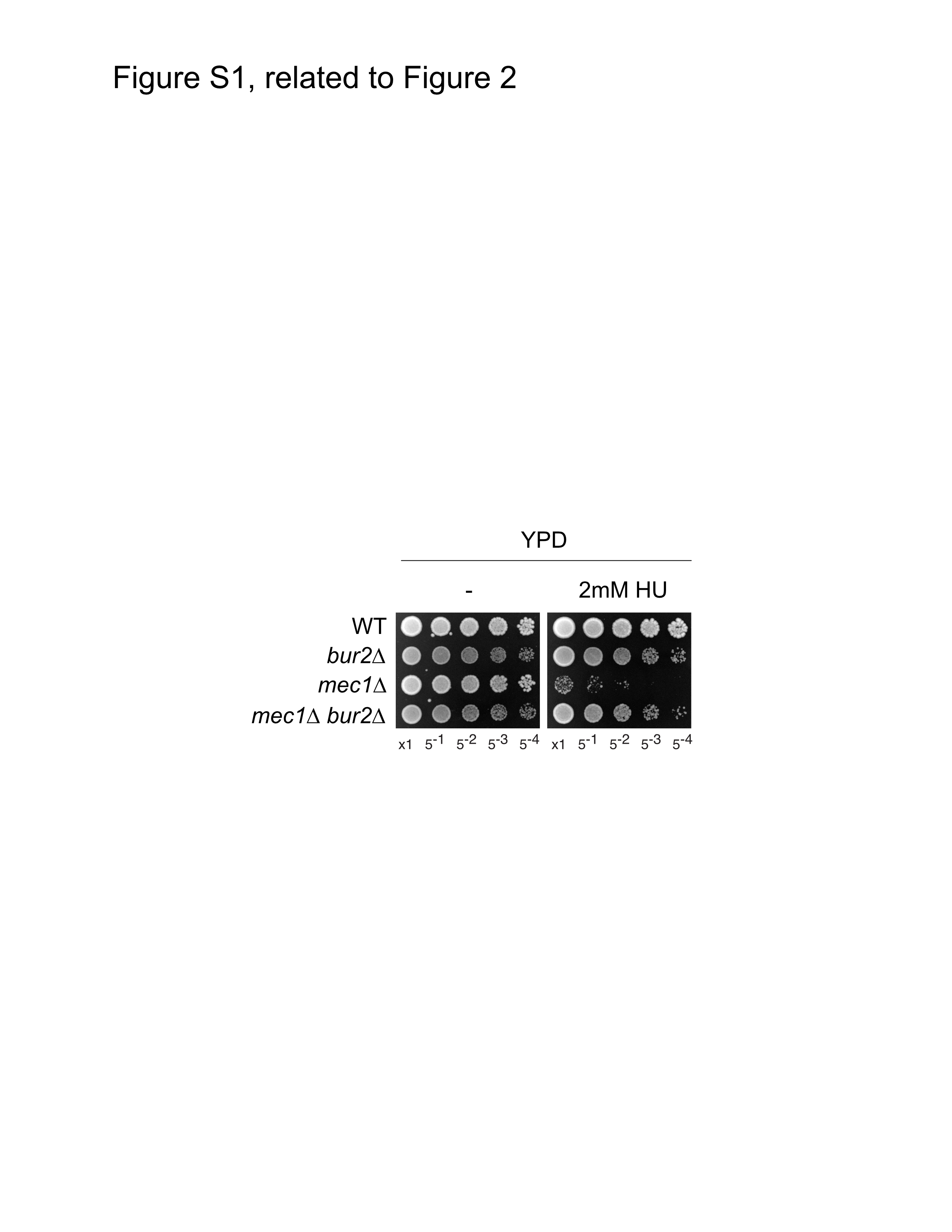

### Supplemental Figure 2

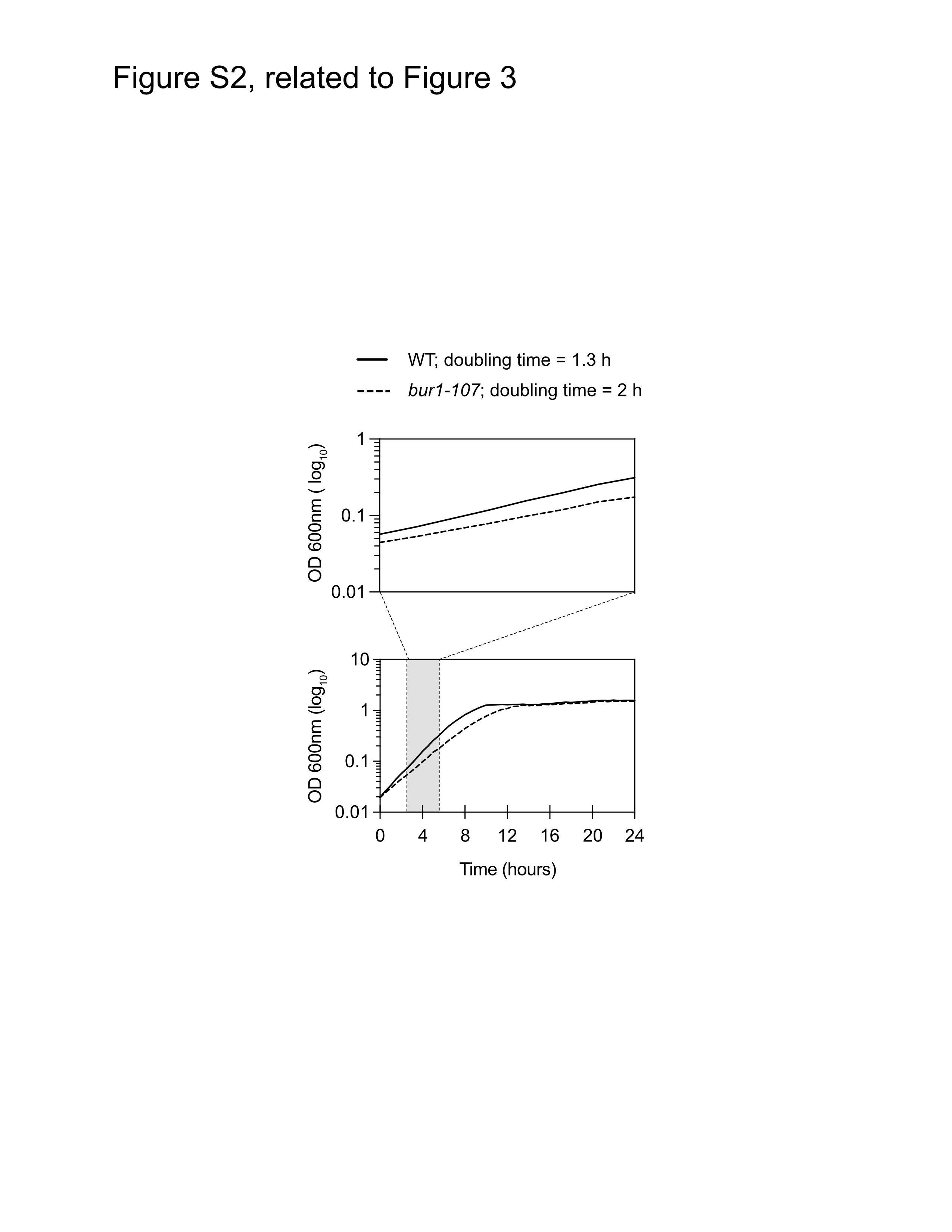

### Supplemental Figure 3

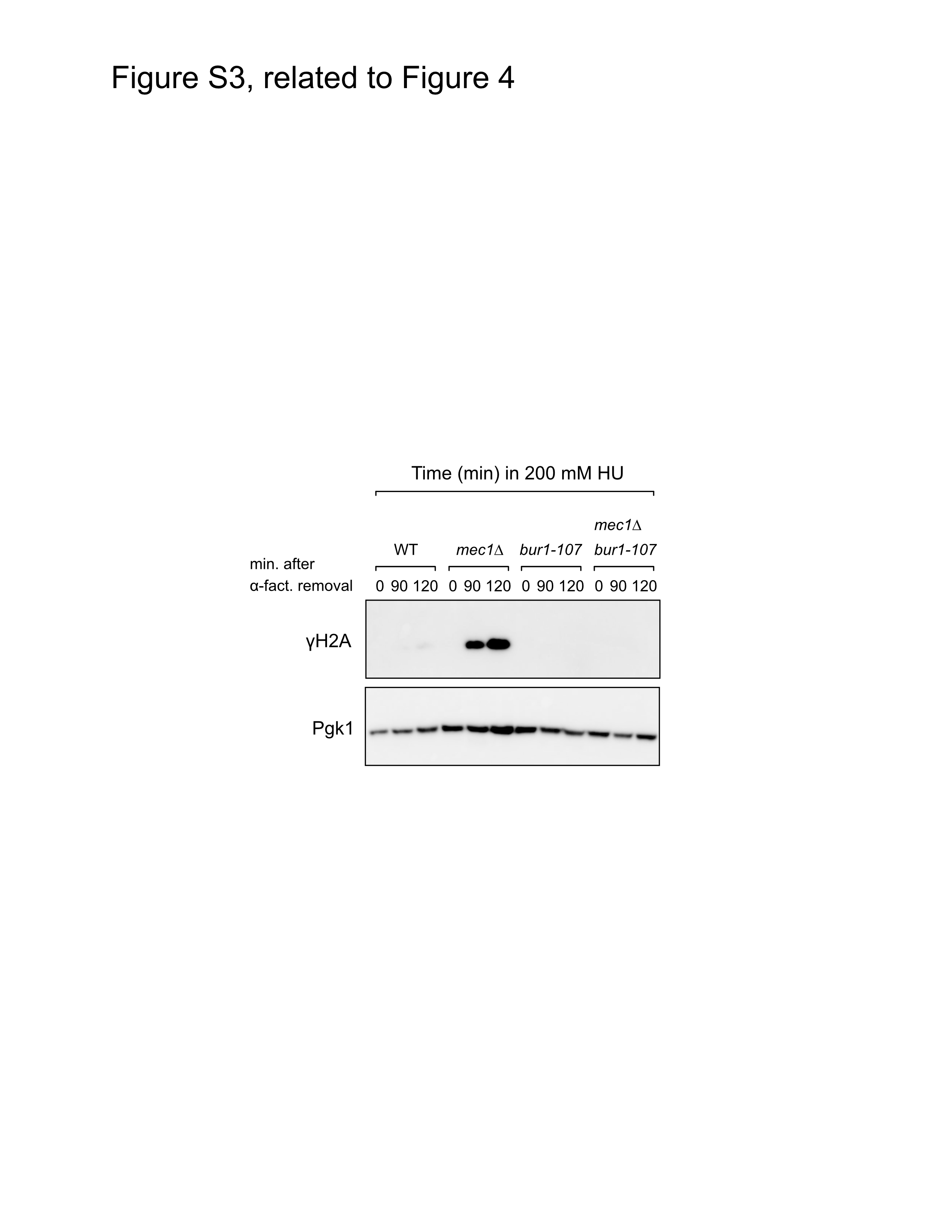

### Supplemental Figure 4

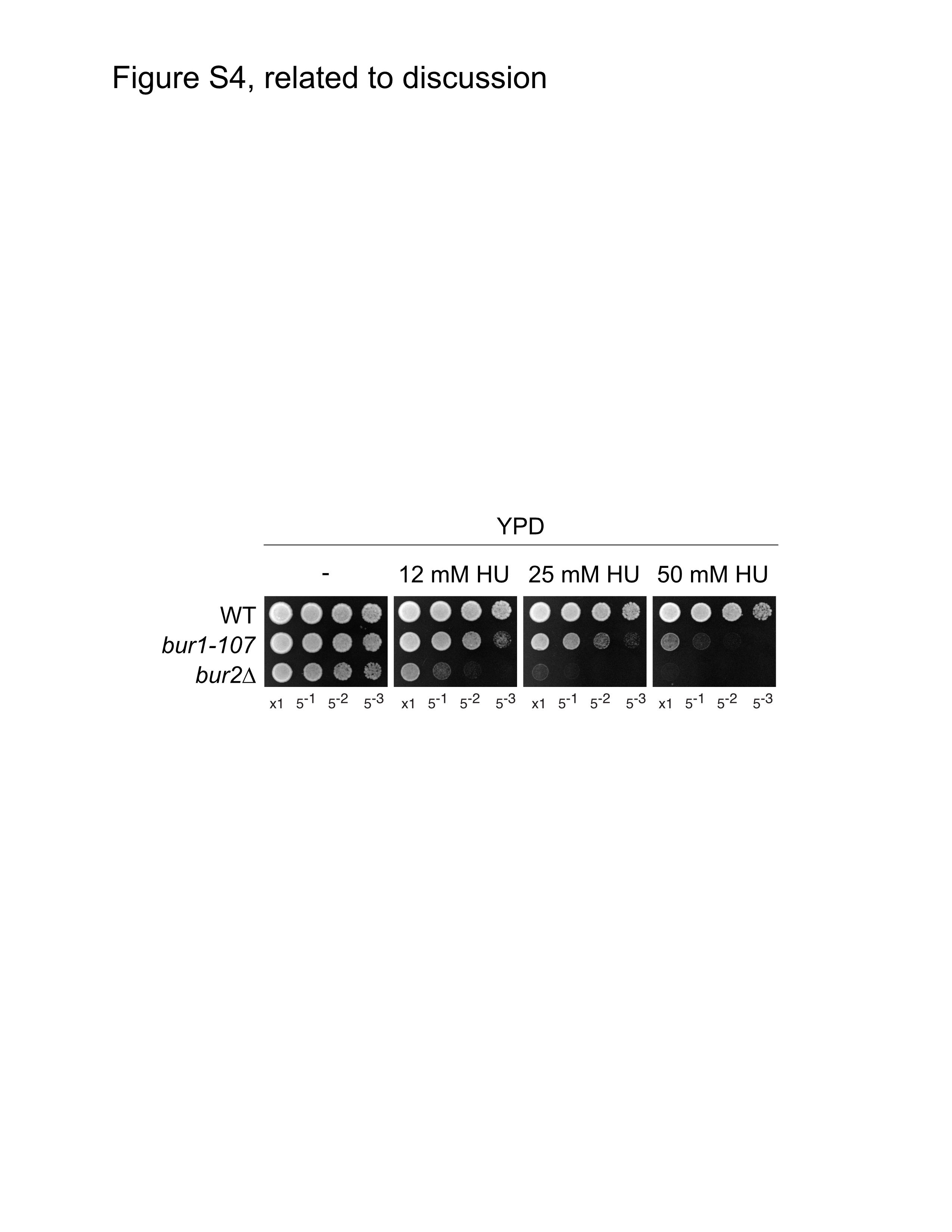
