## Supplemental Table 1 for "Bur1-driven G1-to-S phase transition induces hydroxyurea sensitivity in yeast checkpoint mutants"

| **Code** | **Genotype** | **Background** | **Reference** |
| --- | --- | --- | --- |
| FBO153 | *MATa ura3-52 trp1-63 his3-200 bur1∆::BUR1 KanMX6* | S288C | This study |
| FBO171 | *MATa ura3-52 trp1-63 his3-200 bur1∆::bur1-107 KanMX6* | S288C | This study |
| FBO279 | *MATa ura3-52 trp1-63 his3-200 bur1∆::BUR1 KanMX6 sml1::∆URA3* | S288C | This study |
| FBO283 | *MATa ura3-52 trp1-63 his3-200 bur1∆::bur1-107 KanMX6 sml1∆::URA3* | S288C | This study |
| FBO298 | *MATa ura3-52 trp1-63 his3-200 bur1∆::BUR1 KanMX6 sml1∆::TRP1 mec1∆::HIS3* | S288C | This study |
| FBO304 | *MATa ura3-52 trp1-63 his3-200 bur1∆::bur1-107 KanMX6 sml1∆::TRP1 mec1∆::HIS3* | S288C | This study |
| FBO330 | *MATa ura3-52 trp1-63 his3-200 bur1∆::BUR1 KanMX6 sml1∆::TRP1 rad53∆::HIS3* | S288C | This study |
| FBO338 | *MATa ura3-52 trp1-63 his3-200 bur1∆::bur1-107 KanMX6 sml1∆::TRP1 rad53∆::HIS3* | S288C | This study |
| FBO359 | *MATa ura3-52 trp1-63 his3-200 bur1∆::BUR1 KanMX6 sml1∆::TRP1 mec1∆::HIS3 bar1∆::URA3* | S288C | This study |
| FBO364 | *MATa ura3-52 trp1-63 his3-200 bur1∆::bur1-107 KanMX6 sml1∆::TRP1 mec1∆::HIS3 bar1∆::URA3* | S288C | This study |
| FBO369 | *MATa ura3-52 trp1-63 his3-200 bar1∆::TRP1 bur1∆::BUR1 KanMX6 sml1∆::URA3* | S288C | This study |
| FBO399 | *MATa ura3-52 trp1-63 his3-200 bur1∆::bur1-107 KanMX6 sml1∆::TRP1 bar1∆::URA3* | S288C | This study |
| FBO402 | *MATa ura3-52 trp1-63 his3-200 bur1∆::bur1-107 KanMX6 sml1∆::TRP1 rad53∆::HIS3 bar1∆::NatMX6* | S288C | This study |
| FBO404 | *MATa ura3-52 trp1-63 his3-200 bur1∆::BUR1 KanMX6 sml1∆::TRP1 rad53∆::HIS3 bar1∆::NatMX6* | S288C | This study |
| FBO416 | *MATa his3Δ1 leu2*Δ*0 met15Δ0 ura3Δ0 bur2∆::KanMX6* | BY4741 | Open Biosystems, GE Dharmacon |
| FBO422 | *MATa his3Δ1 leu2*Δ*0 met15Δ0 ura3Δ0 bur2∆::KanMX6 sml1*Δ*::URA3* | BY4741 | This study |
| FBO424 | *MATa his3Δ1 leu2*Δ*0 met15Δ0 ura3Δ0 bur2∆::KanMX6 sml1Δ::URA3 mec1∆::HIS3* | BY4741 | This study |
| FBO433 | *ura3-52/ura3-52 trp1-63/trp1-63 his3-200/his3-200 BUR1/bur1∆::bur1-107 KanMX6 MEC1/mec1∆::HIS3* | S288C | This study |
| FBO547 | *ura3-52/ura3-52 trp1-63/trp1-63 his3-200/his3-200 BUR1/bur1∆::bur1-107 KanMX6 RAD53/rad53∆::HIS3* | S288C | This study |

**Table S1. *Saccharomyces cerevisiae* strains used in this study.**
