## Supplemental Table 2 for "Bur1-driven G1-to-S phase transition induces hydroxyurea sensitivity in yeast checkpoint mutants"

| **Code** | **Vector** | **Purpose** | **Reference** |
| --- | --- | --- | --- |
| pFBO2 | pFA6a-kanMX6 | for gene disruption (KanMX6 marker) | (Bähler et al. 1998) |
| pFBO5 | pRS303 | for gene disruption (HIS3 marker) | (Sikorski and Hieter 1989) |
| pFBO6 | pRS306 | for gene disruption (URA3 marker) | (Sikorski and Hieter 1989) |
| pFBO7 | pRS304 | for gene disruption (TRP1 marker) | (Sikorski and Hieter 1989) |
| pFBO30 | pFA6a-NatMX6 | for gene disruption (NatMX6 marker) | (Hentges et al. 2005) |
| pFBO41 | pFA6a-KanMX6 *BUR1* | for cloning | This study |
| pFBO45 | pFA6a-KanMX6 *bur1-107* | for cloning | This study |
| pFBO112 | pRS316 | for selective growth in SC-URA medium | (Sikorski and Hieter 1989) |

**Table S2. Plasmids used in this study.**

**Literature cited**

Bähler J, Wu J-Q, Longtine MS, Shah NG, Mckenzie III A, Steever AB, Wach A, Philippsen P, Pringle JR. 1998. Heterologous modules for efficient and versatile PCR-based gene targeting in *Schizosaccharomyces pombe*. Yeast. 14(10):943–951. doi:10.1002/(SICI)1097-0061(199807)14:10<943::AID-YEA292>3.0.CO;2-Y.

Hentges P, Van Driessche B, Tafforeau L, Vandenhaute J, Carr AM. 2005. Three novel antibiotic marker cassettes for gene disruption and marker switching in *Schizosaccharomyces pombe*. Yeast. 22(13):1013–1019. doi:10.1002/yea.1291.

Sikorski RS, Hieter P. 1989. A system of shuttle vectors and yeast host strains designed for efficient manipulation of DNA in *Saccharomyces cerevisiae*. Genetics. 122(1):19–27. doi:10.1093/genetics/122.1.19.
